# Multimodal antioxidant and anti-inflammatory activity of *Rauwolfia serpentina* root extracts in experimental models

**DOI:** 10.64898/2026.02.14.705924

**Authors:** Satyabrata Acharya, Smruti Ranjan Das, Aditya Bhaskar Ankari, Sachidanand Nayak

**Affiliations:** Department of Plant Sciences, School of Life Sciences, University of Hyderabad, Hyderabad-500046, Telangana, India

**Keywords:** *Rauwolfia serpentina*, traditional medicine, inflammation, oxidative stress, THP-1 cells, reserpine, NF-κB, IKKα

## Abstract

**Background:** Chronic inflammation and oxidative stress are central drivers of cardiovascular disease progression and remain incompletely addressed by existing pharmacological strategies. Traditional medicinal plants provide a valuable source of multi-target bioactive compounds that may modulate these interconnected pathways. *Rauwolfia serpentina*, a classical antihypertensive plant in Ayurveda, has been historically valued for cardiovascular indications. Yet, its antioxidant and anti-inflammatory actions beyond blood pressure regulation remain insufficiently characterised in immune-driven inflammatory models.

**Methods:** Root extracts of *R. serpentina* prepared using hot and cold ethanol and water were evaluated for antioxidant capacity using DPPH radical scavenging and phosphomolybdenum assays, along with phenolic, flavonoid, and terpenoid quantification. Protective effects against lipid peroxidation were assessed in rat liver and heart homogenates. Anti-inflammatory activity was examined in THP-1 human monocytic cells exposed to lipopolysaccharide (LPS), arachidonic acid (AA), and oxidative stress. Cytokine secretion and gene expression of TNF-α, MCP-1, IL-6, and IL-8 were measured by ELISA and qRT-PCR. Intracellular reactive oxygen species and catalase activity were quantified to assess oxidative regulation. LC-MS-based metabolomic profiling was performed to characterise chemical diversity. The principal alkaloid, reserpine, was evaluated separately, and molecular docking was performed to examine its interaction with IKKα.

**Results:** Ethanolic extracts of *R. serpentina*’s root, particularly the cold ethanolic fraction, showed superior antioxidant capacity, higher phenolic and flavonoid content, and potent inhibition of lipid peroxidation. These extracts markedly suppressed LPS-induced cytokine release and gene expression in THP-1 cells, with pronounced effects on MCP-1 and IL-6. Oxidative stress induced by arachidonic acid was attenuated through reduced intracellular ROS and preservation of catalase activity. Reserpine reproduced key features of the extract response, demonstrating strong suppression of IL-6 and MCP-1 at both transcriptional and secretory levels. Docking analysis indicated stable binding of reserpine within the IKKα catalytic pocket, supporting a plausible mechanism for modulation of the NF-κB pathway.

**Conclusion:** *R. serpentina* root extracts exhibit coordinated antioxidant and anti-inflammatory activity in immune cell models relevant to cardiovascular inflammation. These effects are extraction-dependent and are partially mediated by reserpine through modulation of oxidative stress and inflammatory signalling pathways. The findings support the translational relevance of *R. serpentina* as a traditional medicine with mechanistic activity extending beyond antihypertensive action.

## 1. Introduction

Cardiovascular diseases remain the leading cause of mortality worldwide, driven not only by acute clinical events such as myocardial infarction and stroke but also by progressive vascular damage sustained by chronic inflammation and oxidative stress (Libby, 2021a; Ridker, 2016). Contemporary understanding now recognises cardiovascular pathology as a complex immunometabolic disorder rather than a simple consequence of lipid imbalance or hypertension. Persistent activation of innate immune pathways, excessive reactive oxygen species generation, endothelial dysfunction, and dysregulated cytokine signalling collectively accelerate vascular remodelling and plaque instability (Gimbrone and García-Cardeña, 2016). Pro-inflammatory mediators, including tumour necrosis factor alpha (TNF-α), interleukin 6 (IL-6), and monocyte chemoattractant protein 1 (MCP-1), amplify oxidative injury and promote immune cell recruitment into vascular tissues, reinforcing a self-perpetuating cycle of inflammation and tissue damage (Libby, 2002).

Although current pharmacological strategies have significantly improved survival, many patients continue to exhibit residual inflammatory risk, treatment-associated adverse effects, and incomplete vascular protection (Ridker et al., 2017). These limitations highlight the need for complementary therapeutic approaches that can target inflammatory and oxidative pathways in a broader, potentially safer manner.

Medicinal plants offer a valuable reservoir of bioactive molecules shaped through centuries of empirical use. Among these, *Rauwolfia serpentina* (L.) Benth. ex Kurz (Apocynaceae), commonly known as Sarpagandha or Indian snakeroot, holds a distinctive place in cardiovascular medicine. Traditionally prescribed in Ayurveda for hypertension, anxiety, insomnia, and neurovascular disorders, *R. serpentina* represents one of the earliest examples where a botanical remedy informed modern antihypertensive therapy (Vakil, 1949). Its clinical relevance was historically established through reserpine, an indole alkaloid that revolutionised blood pressure management in the mid-twentieth century.

Beyond reserpine, phytochemical studies reveal that *R. serpentina* roots contain a diverse spectrum of indole alkaloids, including ajmaline, ajmalicine, serpentine, and sarpagine, as well as non-alkaloidal constituents such as phenolics, flavonoids, and terpenoids (Itoh et al., 2005; Kumar et al., 2022). While reserpine is classically associated with monoamine depletion, emerging evidence suggests that *R. serpentina* preparations may exert broader biological effects, including modulation of inflammatory signalling, attenuation of oxidative stress, and stabilisation of endothelial function. These activities are particularly relevant in the context of cardiovascular disease, where macrophage activation and cytokine release play central roles in disease initiation and progression (Libby, 2021a).

Despite its historical importance, systematic evaluation of *R. serpentina* using contemporary cellular and biochemical models of inflammation and oxidative injury remains limited. Most existing studies focus on isolated compounds or blood pressure endpoints, leaving significant gaps in understanding how whole root extracts influence immune-driven vascular pathology. Moreover, mechanistic insight into potential molecular targets linking *R. serpentina* metabolites to inflammatory pathways is still emerging.

In this study, we investigated the antioxidant and anti-inflammatory potential of *R. serpentina* dry root extracts prepared under different extraction conditions, alongside the marker alkaloid reserpine. Using integrated biochemical assays, metabolomic profiling, immune cell-based inflammatory models, and molecular docking, we aimed to characterise how *R. serpentina* modulates oxidative stress, cytokine signalling, and key inflammatory mediators. By combining experimental and in silico approaches, this work seeks to strengthen the mechanistic foundation of *R. serpentina* as a natural product lead for inflammation-driven cardiovascular complications and to support its translational relevance in contemporary integrative medicine.

## 2. Materials and Methods

### Materials

All chemicals were of analytical grade. The following reagents were obtained from Sigma-Aldrich (USA): DPPH, H DCFDA, aluminium chloride (AlCl), arachidonic acid, ascorbic acid, chloroform, dipotassium phosphate (K HPO), DEPC-treated water, ethanol, ferric chloride (FeCl), ferrous ammonium sulfate, 4-fluoro-4′-hydroxybenzophenone, gallic acid, hydrogen peroxide (H O), imidazole, isopropanol, monopotassium phosphate (KH PO), lipopolysaccharide (LPS), 4-nitrophenyl octanoate, orlistat, porcine pancreatic lipase, quercetin, sodium nitrite (NaNO), thiobarbituric acid (TBA), titanium oxysulfate (TiSO), trichloroacetic acid (TCA), Triton X-100, trypan blue, and xylenol orange. Hydrochloric acid (HCl), potassium chloride (KCl), sodium bicarbonate (NaHCO), sodium carbonate (Na CO), sodium hydroxide (NaOH), sulfuric acid (H SO), and Tween-20 were procured from Merck (USA). Ammonium molybdate, Folin–Ciocalteu reagent, and disodium hydrogen phosphate (Na HPO) were purchased from SRL (India). ELISA kits were obtained from BD Biosciences (USA). The iScript cDNA synthesis kit and SYBR Green master mix were sourced from Bio-Rad and Applied Biosystems (USA), respectively. Cell culture reagents, including FBS, L-glutamine, penicillin–streptomycin, RPMI-1640, MTT, and TRIzol, were purchased from Invitrogen (USA).

### 2.1. Plant material collection and authentication

Roots of *R.serpentina* were collected from the Tirumala forest region near Tirupati, Andhra Pradesh, India. Botanical authentication was performed by Dr K. Madhav Chetty, Department of Botany, Sri Venkateswara University, and a voucher specimen was deposited (Voucher No. 0568). Ethical approval for the use of rat tissues was obtained from the Institutional Animal Ethics Committee, University of Hyderabad (UH/IAEC/SY/21/03/2024/46). Plant roots were thoroughly washed to remove soil, shade-dried, and ground to make a fine powder. Samples were stored in airtight containers in the dark until extraction.

### 2.2. Preparation of *R. serpentina* root extracts

Four extract types were prepared to capture metabolites across a polarity spectrum: hot ethanolic extract (HEE), cold ethanolic extract (CEE), hot water extract (HWE), and cold water extract (CWE). Ethanol was selected to enrich alkaloids and moderately polar constituents, while aqueous extraction targeted polar metabolites traditionally accessed in decoctions.

Root powder was extracted following a modified protocol based on the findings of Choudhury et al. (2014), with slight optimisations made for the current study. For hot extraction, 1 g of root powder was mixed with 20 mL of either 80% ethanol or distilled water and stirred at 40°C for 6 h. For cold extraction, 1 g powder was suspended in 4 mL solvent and incubated statically at room temperature for 12–16 h. Extracts were centrifuged at 10,000 rpm for 10 min, filtered, vacuum-dried, and reconstituted on a dry-weight equivalent basis prior to assays.

### 2.3. Antioxidant assays

DPPH radical scavenging activity was assessed by incubating extracts with freshly prepared DPPH solution (0.004% in methanol) for 30 min in the dark, followed by absorbance measurement at 517 nm. Ascorbic acid and gallic acid were used as reference standards. IC_50_ values were calculated from concentration-response curves (Kedare and Singh, 2011).

DPPH scavenging was calculated as: **% inhibition = [(A_control − A_sample) / A_control] × 100**

Total antioxidant capacity was determined using the phosphomolybdenum assay. Samples were incubated with reagent (0.6 M H_2_SO_4_, 28 mM sodium phosphate, 4 mM ammonium molybdate) at 95°C for 90 min, and absorbance was recorded at λ695 nm. Results were expressed as mg ascorbic acid equivalents per gram dry weight of extracts (Prieto et al., 1999).

### 2.4. Phytochemical quantification

To quantify total phenolics, the Folin-Ciocalteu reagent was used with gallic acid standards ranging from 0 to 500 µg/mL, following the method of Singleton et al. (1999). Total flavonoids were quantified by aluminium chloride colourimetry using quercetin standards (Das et al., 2022; Ghorai et al., 2012). Total terpenoids were estimated by the chloroform-sulfuric acid method using linalool calibration (Das et al., 2022; Ghorai et al., 2012). All assays were performed in triplicate.

### 2.5. Pancreatic lipase inhibition

Pancreatic lipase inhibition was assessed using p-nitrophenyl octanoate (p-NPO) as a substrate, following a previously described method Chanda et al., 2019. Pancreatic lipase was incubated with extracts, followed by substrate addition, and absorbance was measured at λ 405 nm. Orlistat served as the positive control. Percentage inhibition was calculated relative to the enzyme control.

Lipase inhibition was calculated as: **% lipase inhibition = 100 − [(As / A) × 100]**
whereas “**As”** is the absorbance in the presence of the inhibitor and “**A”** is the absorbance of the enzyme + substrate control. Assays were performed in triplicate.

### 2.6. Lipid peroxidation in rat tissue homogenates

Liver and heart homogenates were prepared in potassium chloride buffer. Lipid peroxidation was induced using ferric chloride and quantified by TBARS assay via malondialdehyde formation at 532 nm. Gallic acid served as a reference compound (Harlalka et al., 2007).

### 2.7. LC-MS-based metabolite profiling

Root extracts of *R. serpentina* were analysed using LC-MS in positive and negative ion modes on a Shimadzu LC-MS 8045 system with a Phenomenex Kinetex C18 column. Data were processed in MZmine, and metabolites were annotated using KEGG with a confidence score >95% (Kulyal et al., 2021; Sendri et al., 2024).

### 2.8. Cell culture and treatments

THP-1 human monocytic cells were cultured in RPMI 1640 supplemented with fetal bovine serum (FBS) and Pen Strep as an antibiotic. Oxidative stress was induced using arachidonic acid (100 µM), and inflammation was triggered with lipopolysaccharide (0.5 µg/mL) (Kokkiripati et al., 2013).

### 2.9. Cell viability and intracellular ROS

The MTT assay was used to assess cytocompatibility (Kokkiripati et al., 2011). ROS generation was measured using H_2_DCFDA fluorescence following AA stimulation (Bass et al., 1983; Kweon et al., 2001).

### 2.10. Catalase activity

Catalase activity was determined using the titanium oxysulfate method previously described in Koepke et al., (2008), under basal and oxidative conditions, with 3-amino-1,2,4-triazole (3AT) used for validation.

### 2.11. Gene expression studies for inflammatory cytokines (RT-qPCR)

THP-1 cells were pre-treated with the extract for 16 h, followed by LPS stimulation (0.5 µg/mL) for 3 h. Total RNA was isolated using TRIzol with chloroform phase separation and isopropanol precipitation, and purity was assessed by NanoDrop (A260/A280). cDNA was synthesised using iScript™ (Bio-Rad). RT-qPCR was performed with SYBR Green chemistry on a StepOnePlus™ system (10 µL reactions; 95°C for 5 min, 40 cycles of 94°C for 15 s and 60°C for 1 min, followed by melt curve analysis). TNF-α, MCP-1, IL-6, and IL-8 expression was normalised to GAPDH and calculated using the 2^-ΔΔCt method.

### 2.12. Quantification of inflammatory cytokines (ELISA)

THP-1 cells were seeded at 5 × 10 cells/mL in 24-well plates and pre-treated for 12 h with extract equivalents (up to 160 dwt mg/mL) or vehicle. Cells were stimulated with LPS (0.5 µg/mL) for 3 h. Supernatants were clarified and stored at −80°C. TNF-α, MCP-1, IL-6, and IL-8 were quantified using BD OptEIA™ ELISA kits according to manufacturer instructions. Absorbance was measured at 450 nm with reference correction at 570 nm.

### 2.13. Molecular docking

Reserpine docking against human IKKα (PDB ID: 5EBZ) was performed using AutoDock Vina. Protein and ligand preparation were done in AutoDockTools. The best-ranked poses were analysed using PyMOL and BIOVIA Discovery Studio Visualiser (Eberhardt et al., 2021; Trott and Olson, 2010).

A molecular docking study was conducted on the human LPS-induced pro-inflammatory factor IKKα (PDB ID: 5EBZ) to evaluate its interaction with the standard inhibitor 5TL (PubChem CID: 16048085) and reserpine (PubChem CID: 5770). The 3D structures of the protein (IKKα) and the ligands (5TL and reserpine) were obtained from the Protein Data Bank (PDB)(https://www.rcsb.org/) and the PubChem database (https://pubchem.ncbi.nlm.nih.gov/), respectively. The protein and ligand preparations were done using Autodock4 v4.2.6. The resulting molecular docking-compatible files of the protein and 5TL were subjected to blind rigid docking using AutoDock Vina v1.1.2 (Eberhardt et al., 2021; Trott and Olson, 2010) with 5 different random seed values. The resulting seed value, which provided the best binding affinity between protein and 5TL, was used for blind rigid docking of protein and reserpine. The Autodock Vina results were visualised and analysed using PyMOL (The PyMOL Molecular Graphics System, Version 3.1, Schrödinger, LLC) and Biovia Discovery Studio Visualizer version 2025 (Biovia, D.S. Discovery Studio Visualizer 2025, San Diego)

### 2.14. Statistical analysis

All experiments were performed in biological triplicates unless stated otherwise. Results are expressed as mean ± standard deviation. Statistical comparisons were conducted using one-way ANOVA. A p-value less than 0.05 was considered statistically significant.

## 3. Results

### 3.1. Antioxidant activity of *Rauwolfia serpentina* root extracts

The DPPH assay was employed to evaluate the free-radical scavenging activity of *R. serpentina* root extracts obtained under various extraction conditions. Gallic acid, used as a reference antioxidant, showed an IC of 2.64 µg/mL (Supplementary Figure 1), validating assay performance. Among all extracts, CEE demonstrated the strongest DPPH scavenging activity, with an IC of 33.84 ± 2.92 µg/mL, indicating superior free radical neutralisation capacity. This was followed by the HEE (IC = 55.00 ± 6.90 µg/mL). In contrast, both aqueous preparations were less effective, with IC values of 60.14 ± 1.85 µg/mL for HWE and 66.22 ± 5.20 µg/mL for CWE (Supplementary Figure 2). Reserpine displayed moderate antioxidant activity (IC = 44.68 µM), indicating that the pronounced antioxidant effects of CEE likely arise from synergistic contributions of multiple metabolites rather than reserpine alone (Supplementary Figure 3).

Total antioxidant capacity assessed by the phosphomolybdenum method revealed a distinct pattern. HWE exhibited the highest reducing power (16.86 ± 0.29 mg GAE/g dw), whereas HEE, CEE, and CWE showed comparable values ranging from 10.36 to 10.81 mg GAE/g dry weight (Supplementary Figure 4). This divergence suggests that heating under aqueous conditions favours the extraction of bulk reducing compounds, while ethanol preferentially enriches radical-scavenging constituents.

### 3.2. Phytochemical composition of root extracts

Quantification of major phytochemical classes demonstrated marked enrichment in ethanol-derived extracts. CEE contained the highest total phenolic content (67.66 ± 3.08 mg GAE/g dw) and flavonoid content (89.93 ± 1.95 mg QE/g dry weight), consistent with its superior DPPH scavenging profile. HEE showed moderate phenolic levels (24.07 ± 0.78 mg GAE/g dry weight), while aqueous extracts contained substantially lower phenolics and flavonoids (Supplementary Figure 5A-B).

In contrast, terpenoid content was maximal in HEE (2.78 ± 0.03 mg LE/g dry weight), whereas HWE and CWE showed minimal terpenoid enrichment (Supplementary Figure 5C). These extraction-dependent phytochemical patterns provide a biochemical basis for the differential antioxidant and anti-inflammatory activities observed across extracts.

### 3.3. Pancreatic lipase inhibition

Pancreatic lipase inhibition was evaluated to assess whether *Rauwolfia serpentina* root extracts could modulate lipid digestion, a process closely linked to metabolic inflammation and cardiovascular risk. Orlistat, used as the positive control, reduced enzyme activity to 45.00%, confirming assay sensitivity.

Among the extracts, the cold extracts displayed the strongest inhibitory activity. CEE reduced lipase activity from 61.95 ± 1.84% at a concentration of 80 µg/mL to 48.07 ± 2.15% at 120 µg/mL, reaching 40.10 ± 0.77% at 160 µg/mL. A comparable pattern was observed for CWE, which produced enzyme activity values of 61.95 ± 1.65%, 48.07 ± 2.28%, and 40.10 ± 0.92% across the same concentration range. These results indicate strong dose-dependent inhibition, approaching the efficacy of orlistat at the highest dose. HEE showed modest suppression, decreasing activity from 96.99 ± 1.59% at 80 µg/mL to 70.20 ± 1.35% at 160 µg/mL. In contrast, HWE exhibited a delayed but sharp inhibitory response at the highest concentration, with enzyme activity dropping to 41.13 ± 0.74% (Figure 1 A-D).

**Figure 1.**
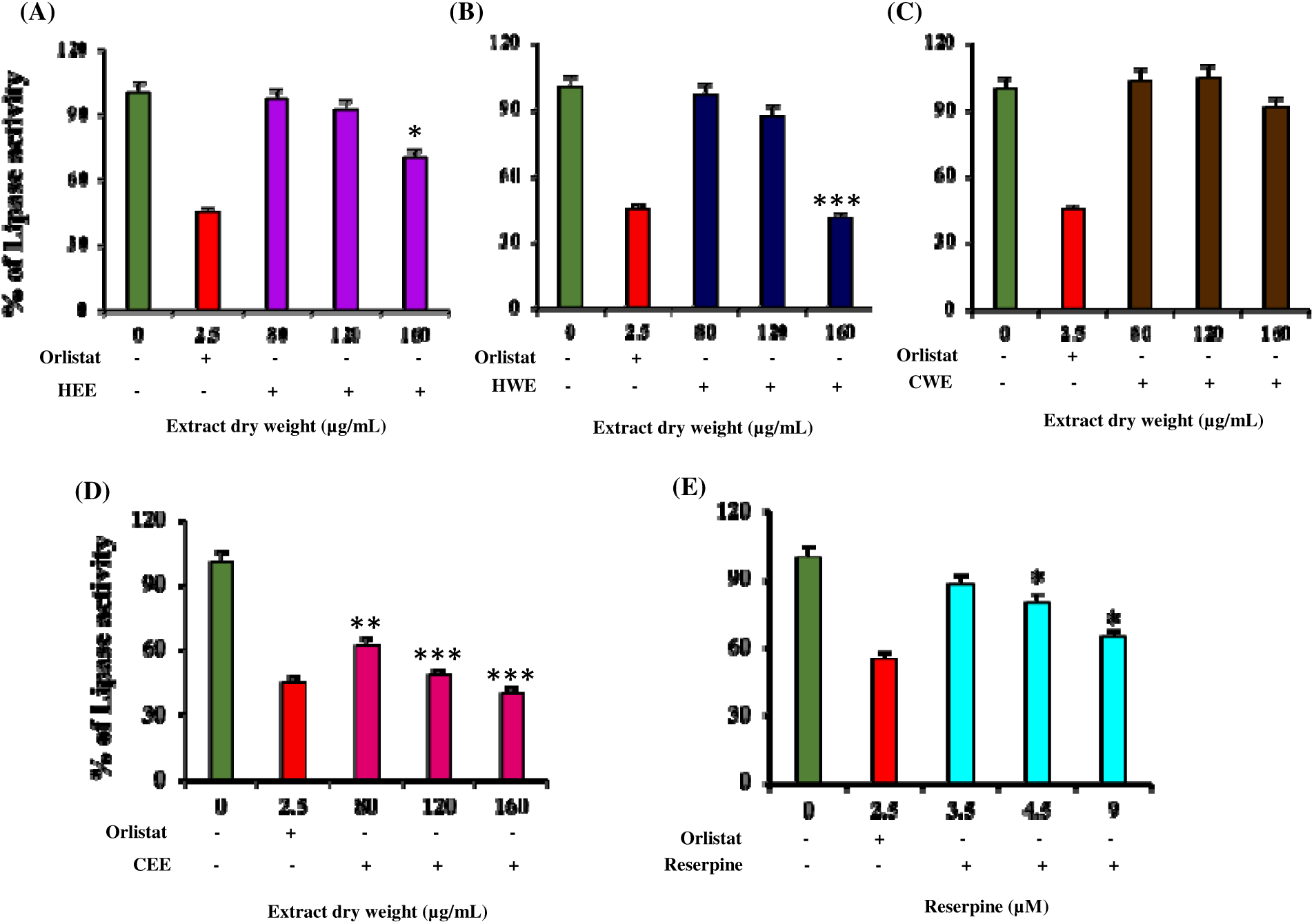
Pancreatic lipase inhibitory activity of *Rauwolfia serpentina* root extracts and reserpine. Dose-dependent inhibition of pancreatic lipase by (A) HEE, (B) HWE, (C) CWE, (D) CEE, and reserpine. Data are presented as mean ± standard deviation from three independent experiments (n = 3). All treatment groups exhibited statistically significant inhibition compared with the enzyme control (p < 0.001). Orlistat was used as the reference inhibitor.

Reserpine showed comparatively weak inhibition, maintaining enzyme activity above 64% even at the maximal tested concentrations (Figure 1E). Together, these findings suggest that pancreatic lipase inhibition is primarily mediated by non-alkaloidal constituents enriched in cold extracts, likely involving phenolics and flavonoids identified in the metabolomic analysis.

### 3.4. Protection against lipid peroxidation in tissue homogenates

In liver homogenates, ferric chloride induced marked lipid peroxidation, which was significantly attenuated by the extracts. CEE exhibited the strongest protection with an IC of 16.66 ± 1.27 µg/mL, followed by CWE (36.53 ± 0.12 µg/mL). Thermally prepared extracts showed limited activity (HEE: 134.77 ± 0.12 µg/mL; HWE: 154.12 ± 1.22 µg/mL). Reserpine displayed moderate inhibition (IC = 9.89 ± 0.16 µM), supporting partial alkaloid contribution (Figure 2).

**Figure 2.**
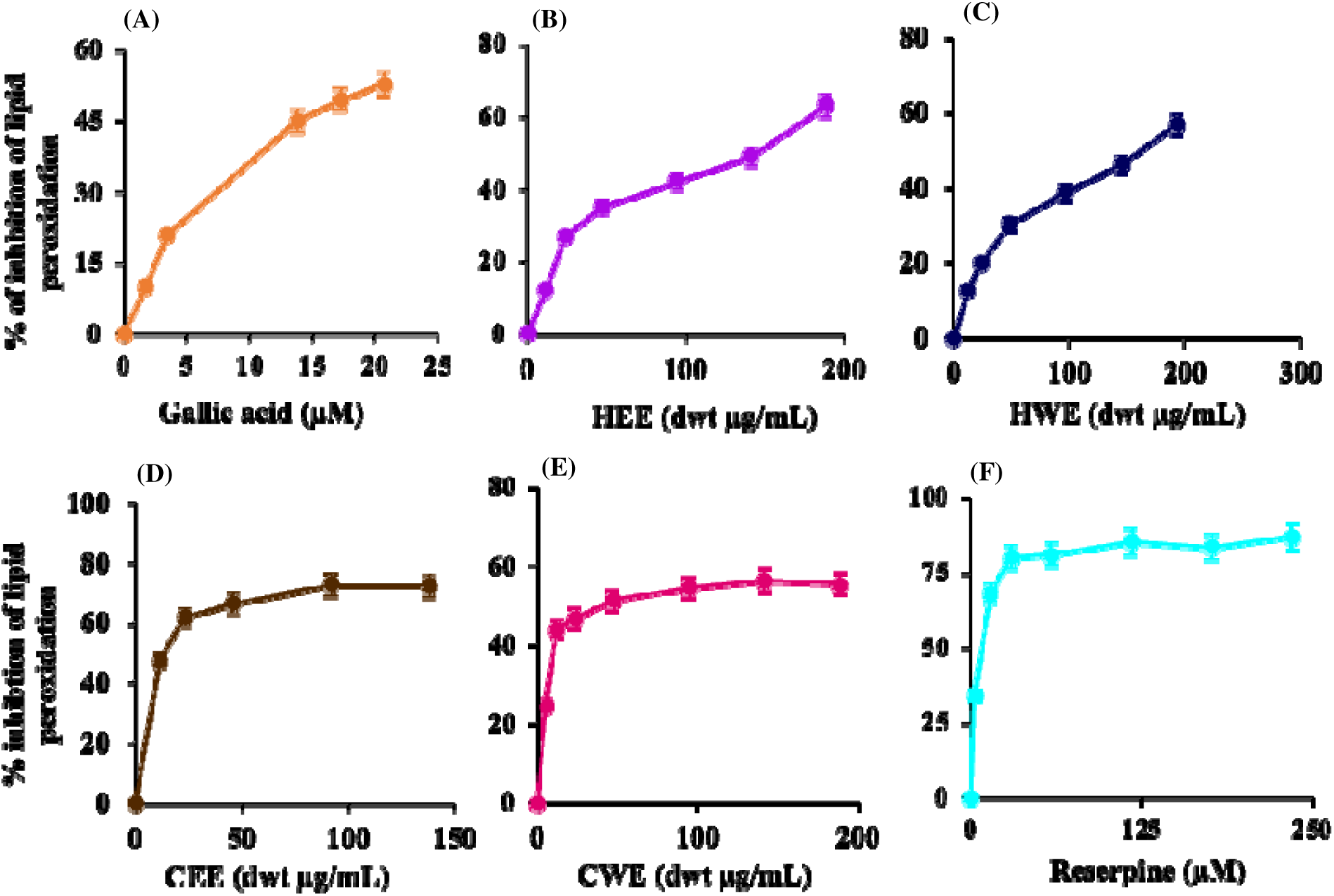
Inhibition of lipid peroxidation by *Rauwolfia serpentina* root extracts and reserpine in rat heart homogenates. Protective effects of gallic acid (A, reference standard), followed by *R. serpentina* dried root extracts (B-E) and reserpine (F) against FeClD-induced lipid peroxidation in rat cardiac tissue, measured by TBARS assay. Data are expressed as mean ± standard deviation from three independent experiments (n = 3). Extracts and reserpine exhibited concentration-dependent suppression of malondialdehyde formation

Cardiac homogenates showed comparatively weaker protection. HEE exhibited the lowest IC (148.76 ± 3.07 µg/mL), followed by CEE (164.73 ± 4.00 µg/mL), while aqueous extracts exceeded 166 µg/mL. Gallic acid showed strong reference inhibition (18.98 ± 0.47 µM), whereas reserpine retained moderate activity (25.08 ± 2.90 µM) (Figure 3).

**Figure 3.**
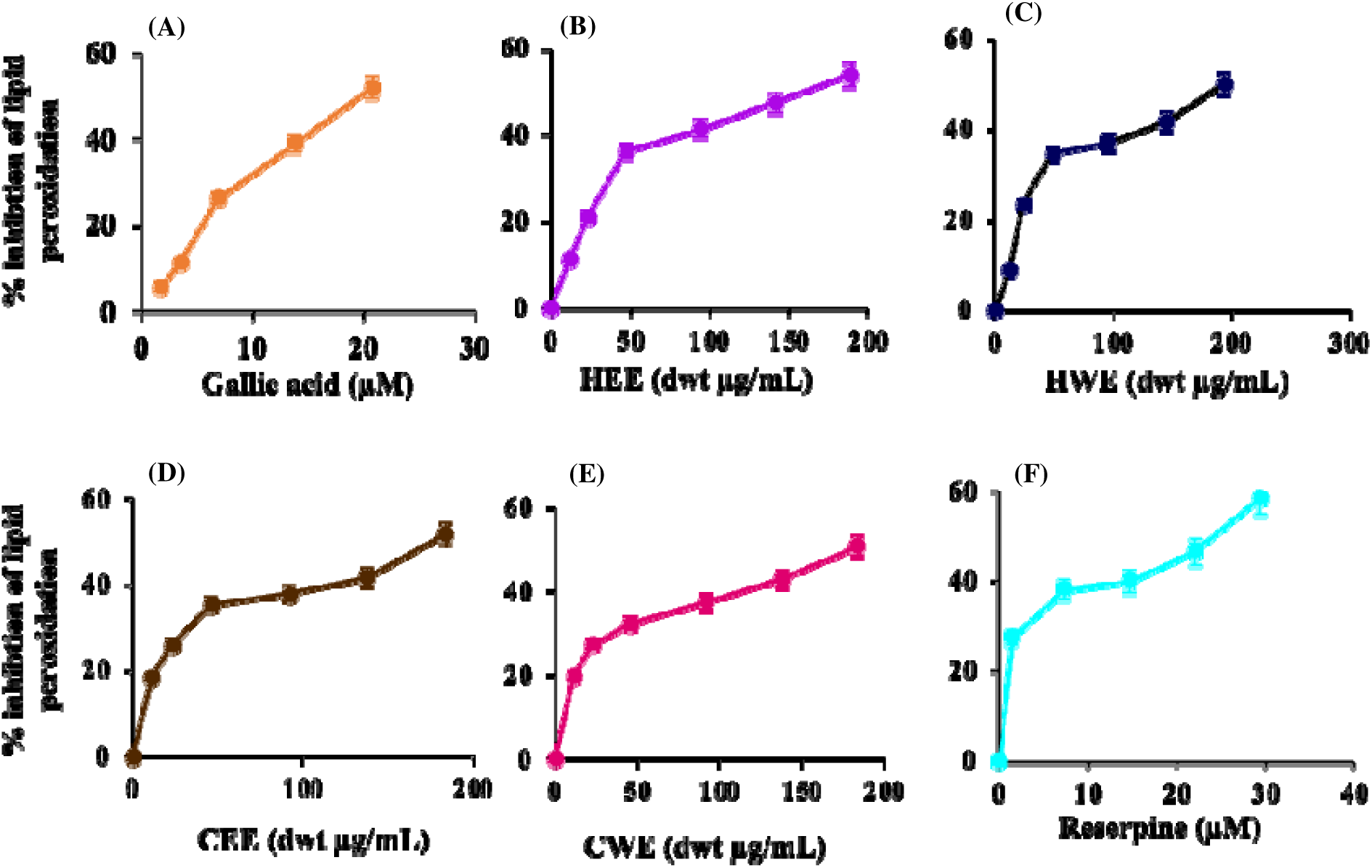
Inhibition of lipid peroxidation by *Rauwolfia serpentina* root extracts and reserpine in liver homogenates. Protective effects of gallic acid (A, reference standard), followed by *R. serpentina* dried root extracts (B-E) and reserpine (F) against FeCllJ-induced lipid peroxidation in rat liver tissue, measured by TBARS assay. Data are expressed as mean ± standard deviation from three independent experiments (n = 3). Extracts and reserpine exhibited concentration-dependent suppression of malondialdehyde formation compared with the oxidative control.

### 3.5. Metabolomic profiling

LC-MS analysis identified 105 metabolites spanning alkaloids, phenolics, lignans, terpenoids, fatty acids, sterols, and quinones. Alkaloids dominated the metabolome, with 73 compounds, including reserpine, ajmaline, serpentine, and sarpagine (Supplementary Table 1). Notably, 21 alkaloids not previously reported in *R. serpentina* were detected (Table 1). Several non-alkaloidal metabolites with known antioxidant or anti-inflammatory potential, such as verbascoside, hinokinin, paradol, and trimethoxybenzoic acid derivatives, were also identified, underscoring the chemical diversity underlying extract bioactivity.

**Table 1.**
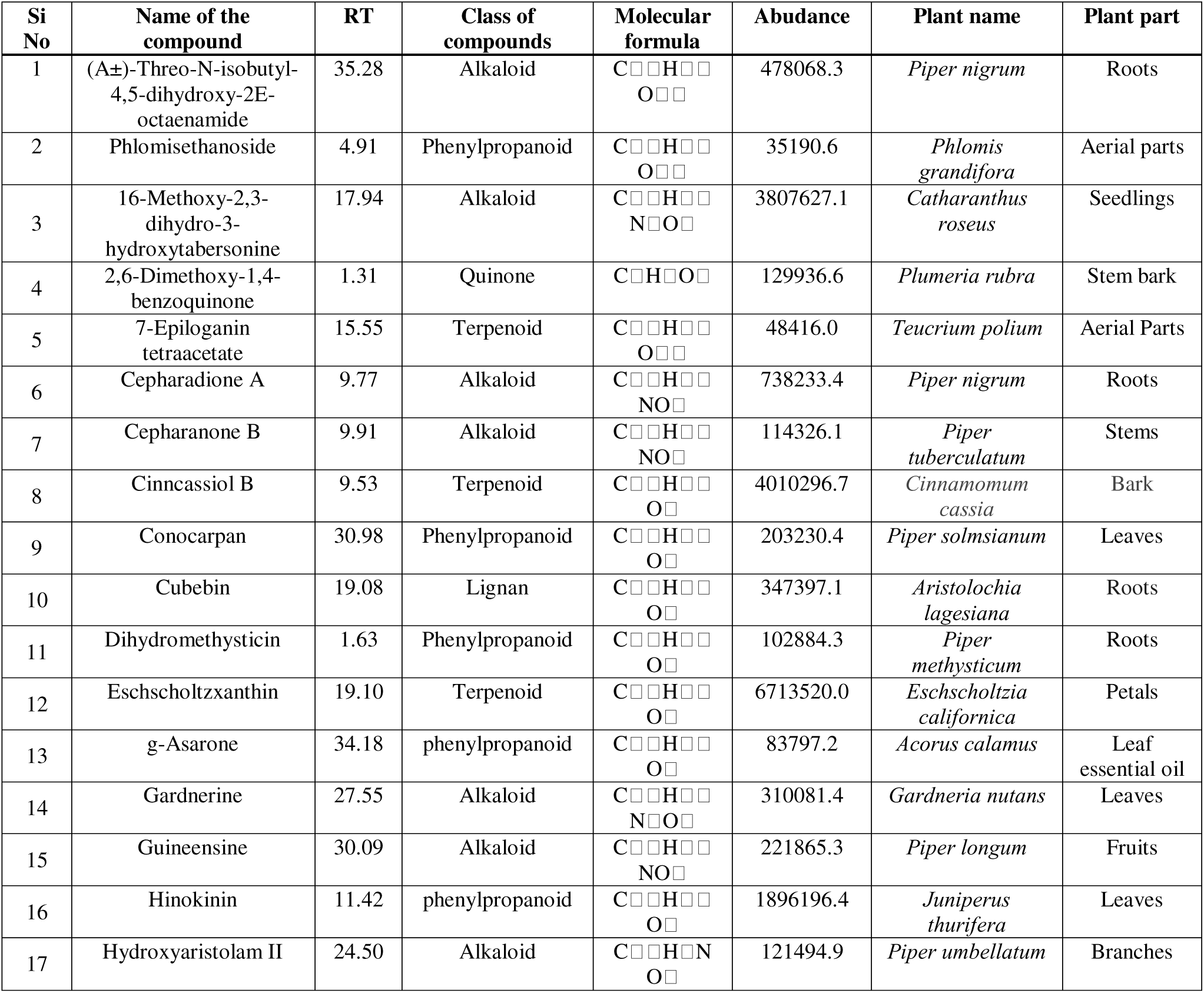
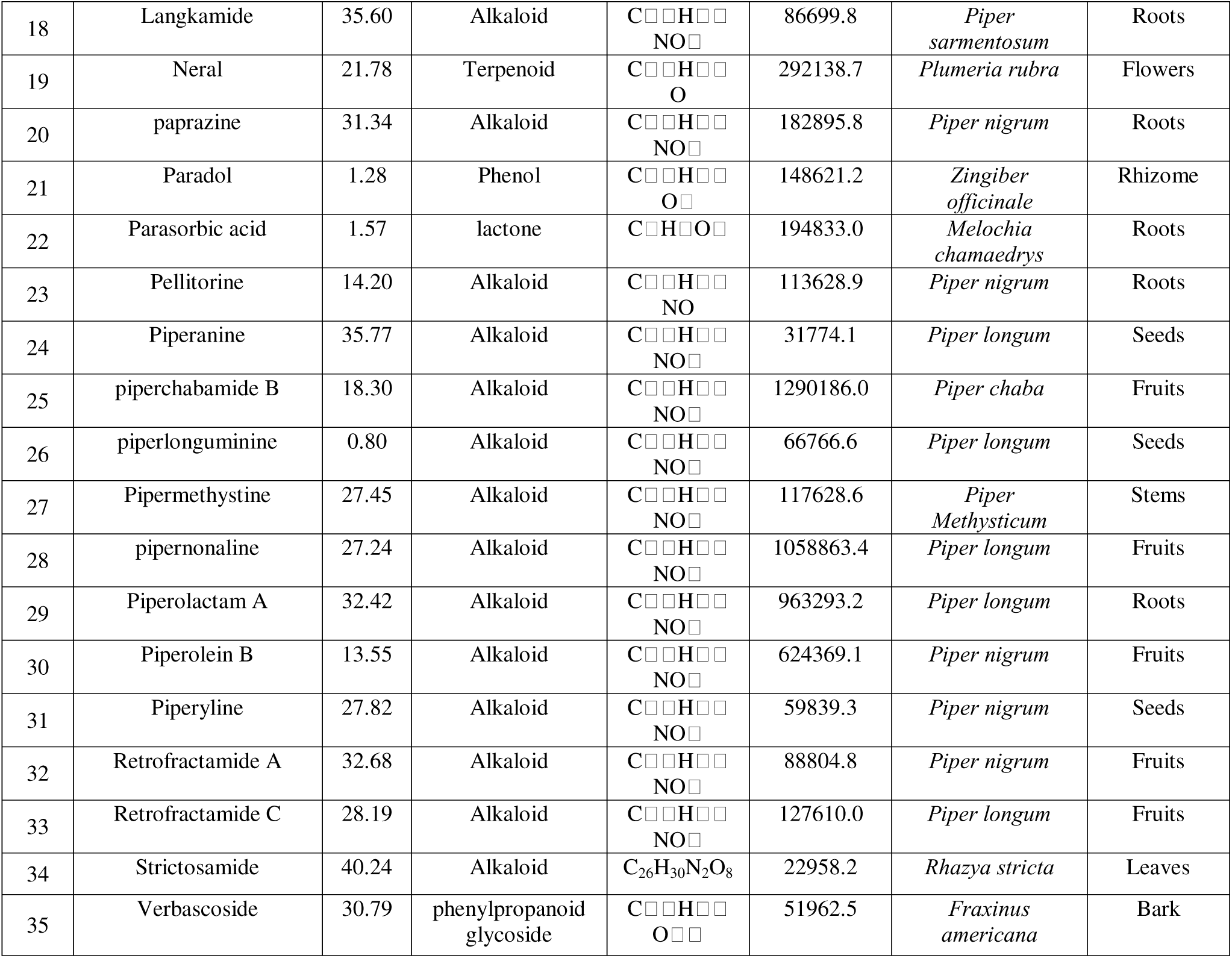
Metabolites tentatively identified for the first time in the LC–MS analysis of dried root extracts of *Rauwolfia serpentina*.

### 3.6. Cytotoxicity and oxidative stress modulation

All the extracts maintained THP-1 cell viability above 90% at concentrations up to 160 µg/mL. Reserpine was well tolerated up to 100 µM, with mild cytotoxicity observed only at higher doses (Supplementary Figure 6).

To evaluate antioxidant activity under inflammatory stress, intracellular ROS generation was measured following arachidonic acid stimulation. In unstimulated cells, ROS levels remained low, confirming that *R. serpentina* extracts did not induce oxidative stress under basal conditions. Arachidonic acid exposure induced a marked increase in ROS, validating the oxidative challenge model.

Among all the extracts, HEE showed the strongest and most consistent suppression of ROS. ROS values decreased from 10.60 ± 0.10 at 40 µg/mL to 10.29 ± 0.16 at 80 µg/mL, then fell sharply to 6.10 ± 0.15 at 120 µg/mL and remained suppressed at 6.35 ± 0.16 at 160 µg/mL. CEE also significantly reduced ROS, lowering values to 8.80 ± 0.11 at 40 µg/mL and maintaining stable suppression across higher concentrations (8.54 ± 0.23 at 120 µg/mL). In contrast, aqueous extracts showed minimal protective effects, with ROS levels remaining close to the arachidonic acid control (Figure 4A-D).

**Figure 4.**
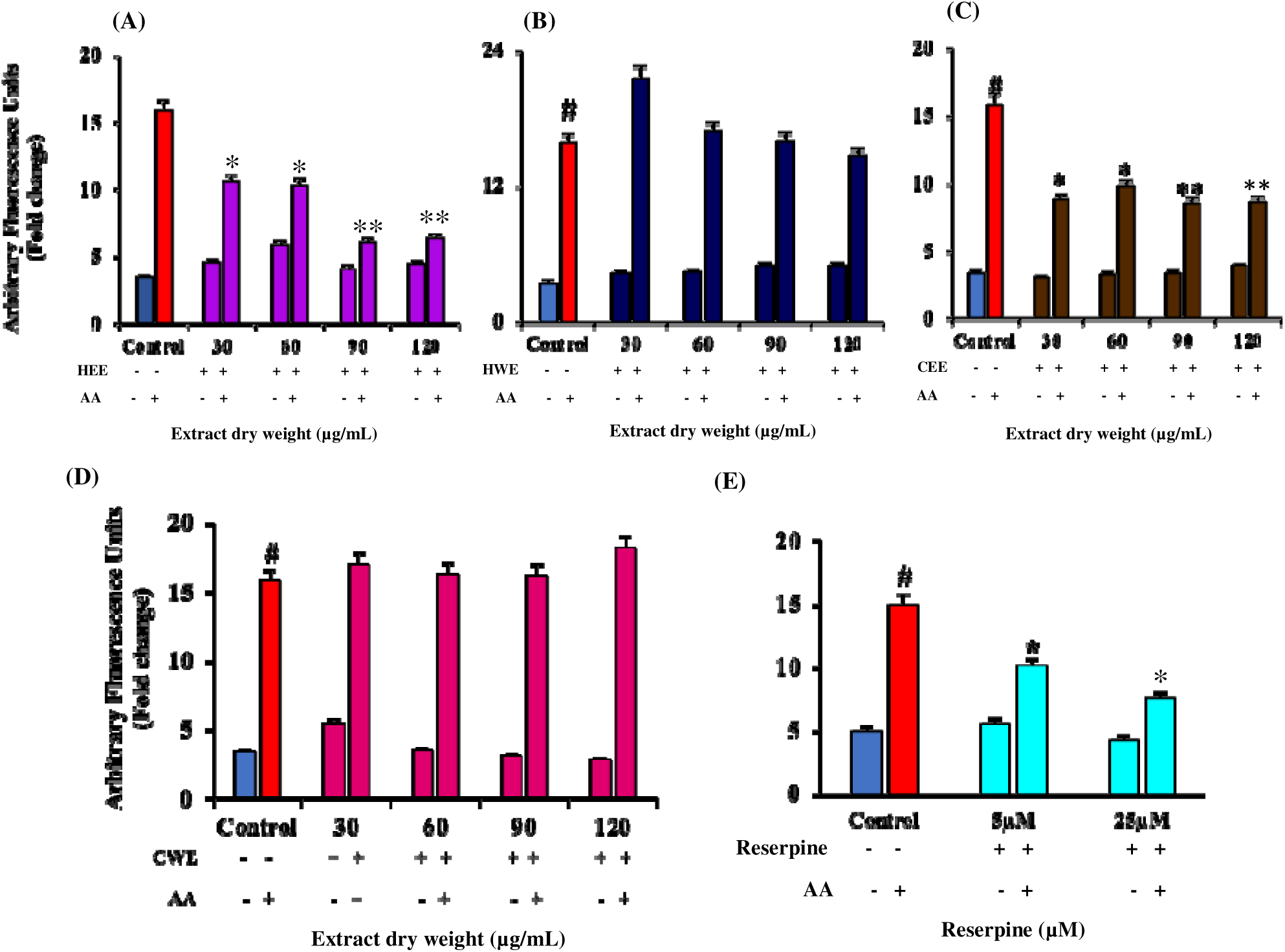
Effect of *Rauwolfia serpentina* root extracts and reserpine on arachidonic acid-induced intracellular ROS generation in THP-1 cells. Cells were treated with (A) HEE, (B) HWE, (C) CEE, (D) CWE, or reserpine (E) prior to arachidonic acid stimulation. ROS levels were measured using HDDCFDA fluorescence. Values are expressed as mean ± SD (n = 3). All treatments showed significant differences compared with the arachidonic acid control (p < 0.001).

At the compound level, reserpine reduced ROS to 10.16 ± 0.19 at 5 µM and further to 7.62 ± 0.11 at 25 µM, demonstrating concentration-dependent antioxidant activity (Figure 4E). These data indicate that ethanol-soluble metabolites, including reserpine, effectively mitigate lipid-mediated oxidative stress in immune cells.

### 3.7. Catalase activity

To assess whether *R. serpentina* extracts boost endogenous antioxidant defence under basal conditions, during arachidonic acid oxidation, and in the presence of 3AT, catalase activity was evaluated. Under basal conditions, catalase activity increased most notably with HEE at 120 µg/mL (123.99 ± 3.41) and CEE at 120 dwtµg/mL (121.37 ± 0.91), while CWE showed weaker activation at 90 µg/mL (71.78 ± 1.90). During AA-induced oxidative stress, HWE maintained catalase near the control range (93.65 ± 1.77 at 90 µg/mL and 95.40 ± 0.66 at 120 µg/mL), whereas CEE showed stronger preservation at higher doses (112.89 ± 5.36 at 120 µg/mL). Under direct catalase inhibition with 3AT, CEE showed the strongest protection, maintaining catalase activity at 97.09 ± 0.70 at 90 µg/mL and 82.30 ± 2.94 at 120 µg/mL, whereas HEE and HWE retained lower activity, near 60-68%. Under combined stress (3AT+AA), catalase retention was highest with CEE at 120 µg/mL (80.93 ± 0.29). CWE also showed a strong dose response, rising to 82.99 ± 4.14 at 120 µg/mL (Supplementary Figure 7A-D).

Reserpine produced a pronounced catalase-enhancing effect. Basal catalase activity increased to 129.76 ± 1.05 at 5 µM and 140.58 ± 0.83 at 25 µM. Under AA challenge, catalase remained elevated at 114.80 ± 0.90 and 117.99 ± 4.25, respectively. Importantly, under combined 3AT and AA stress, reserpine retained catalase activity at 80.20 ± 1.48 at 5 µM and 93.19 ± 6.71 at 25 µM, indicating strong enzyme-level resilience during dual oxidative and inhibitory stress (Supplementary Figure 7E).

### 3.8. Modulation of inflammatory gene expression by *Rauwolfia serpentina* extracts in LPS-stimulated THP-1 cells

THP-1 cells were stimulated with LPS to study the effects of *Rauwolfia serpentina* on inflammatory signalling at the transcriptional level. TNF-α, MCP-1, IL-6, and IL-8 levels were evaluated using qRT-PCR. LPS induced a strong cytokine and chemokine response, setting a baseline for extract-mediated suppression.

LPS markedly upregulated TNF-α to 543.94 ± 27.20-fold. This response was predominantly unchanged with HWE and CWE but exhibited a substantial drop with the ethanolic extracts, reaching 211.49 ± 10.57-fold with HEE and 36.83 ± 1.79-fold with CEE. MCP-1 had a similar pattern. LPS increased MCP-1 to 67.64 ± 3.38-fold, but both ethanolic extracts reduced it to near basal levels, declining to 11.12 ± 0.56-fold with HEE and 4.00 ± 0.16-fold with CEE. In contrast, HWE and CWE demonstrated minimal protection against MCP-1. IL-6 decreased the most with ethanol extracts. Although LPS elevated IL-6 to 418.69 ± 20.93-fold, reduced it to 9.00 ± 0.44-fold and CEE to 15.31 ± 0.77-fold. HWE and CWE had no significant impact. The IL-8 responses showed minimal variation. LPS induced a 375.68 ± 18.78-fold elevation in IL-8 levels. HEE achieved nearly complete suppression, decreasing IL-8 to 5.02 ± 0.25-fold. In contrast, CEE showed little effectiveness for IL-8, and both water extracts showed an inconsequential response to LPS induction (Figure 5).

**Figure 5.**
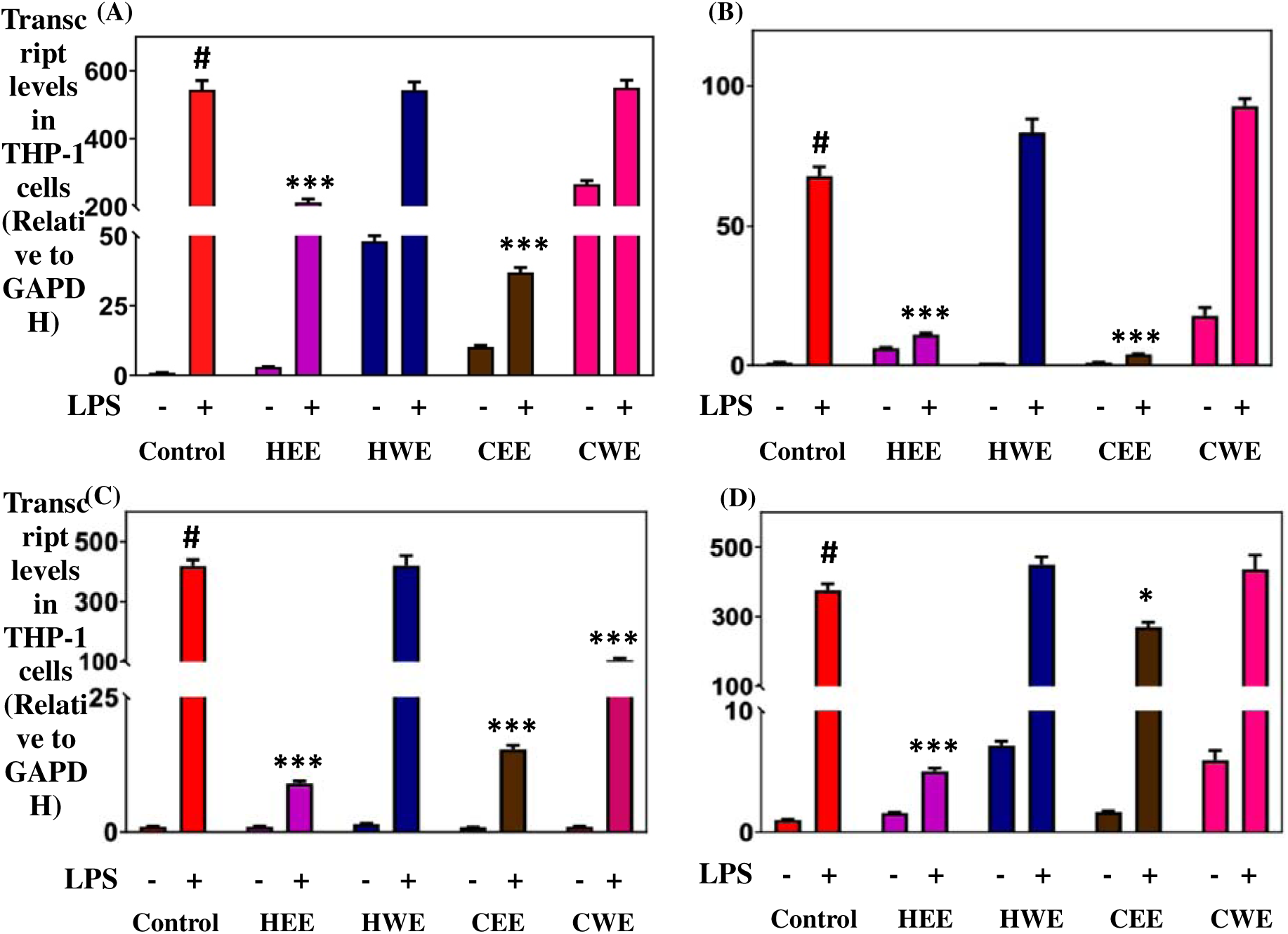
Modulation of pro-inflammatory gene expression by *Rauwolfia serpentina* root extracts in LPS-stimulated THP-1 cells. Cells were pre-treated with *R. serpentina* extracts, followed by LPS stimulation for 3 h. Relative mRNA expression of TNF-α (A), MCP-1 (B), IL-6 (C), and IL-8 (D) was quantified by qRT-PCR using GAPDH as the internal control. Data are presented as mean ± SD from three independent experiments (n = 3). #p < 0.001 vs. untreated control; *p < 0.001 vs. LPS-treated group.

Reserpine significantly reduced LPS-induced inflammatory gene expression in THP-1 cells. TNF-α levels decreased from 20.00 ± 0.88-fold to 18.83 ± 0.94-fold at 5 µM and to 15.89 ± 0.79-fold at 25 µM. MCP-1 expression was strongly suppressed from 78.45 ± 3.92-fold to 3.79 ± 0.19-fold at 5 µM and 4.73 ± 0.24-fold at 25 µM. IL-6 expression declined from 750.00 ± 30.00-fold to 9.87 ± 0.49-fold and 5.36 ± 0.27-fold at 5 µM and 25 µM, respectively. IL-8 expression was also reduced in a concentration-dependent manner, decreasing from 289.96 ± 14.50-fold to 119.26 ± 5.96-fold at 5 µM and 63.16 ± 3.16-fold at 25 µM (Figure 6).

**Figure 6.**
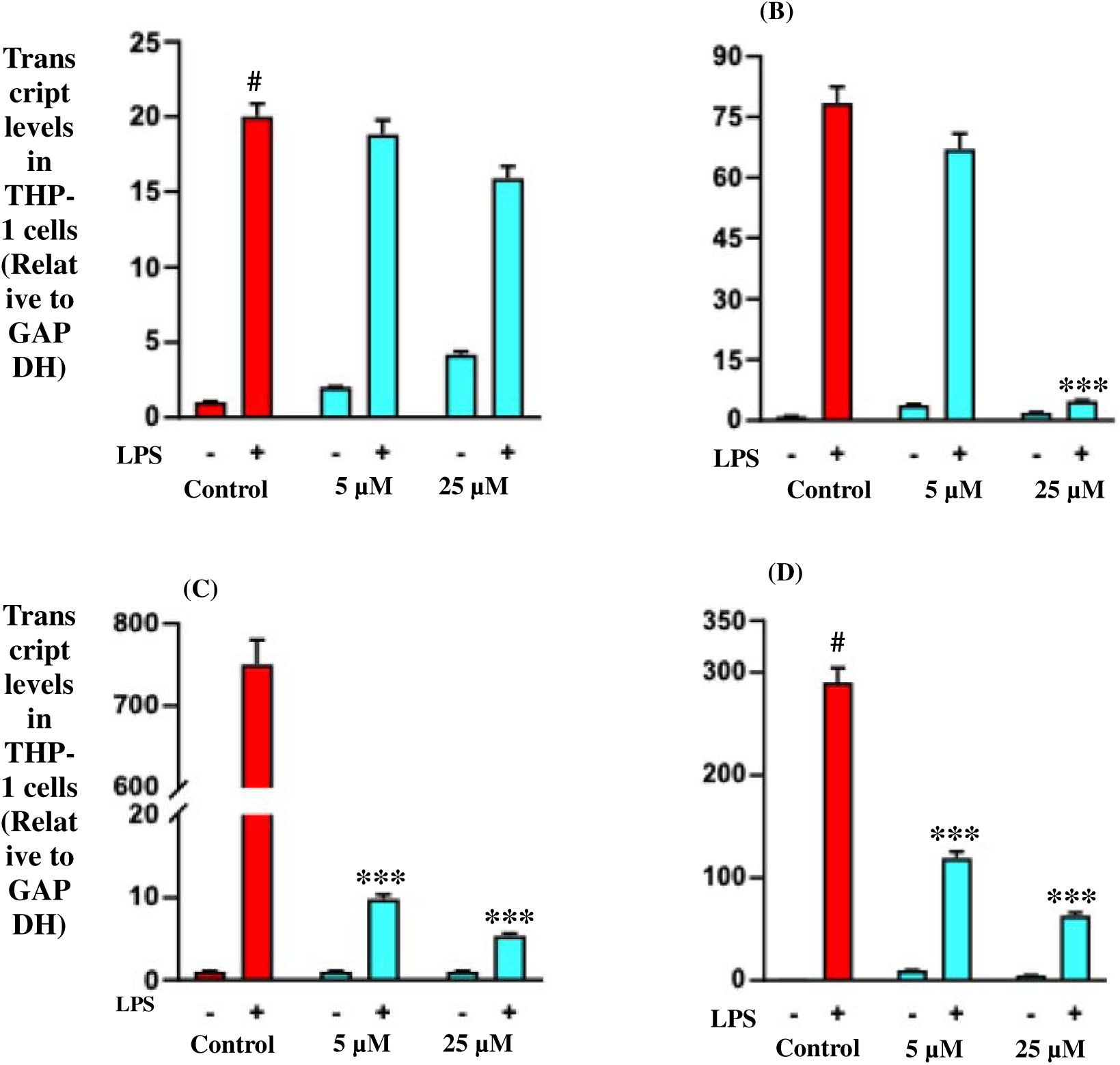
Effect of reserpine on the expression of inflammatory genes induced by LPS. THP-1 cells were exposed to varying doses of reserpine acid before LPS stimulation for 3 hours. Expression levels of TNF-α (A), MCP-1 (B), IL-6 (C), and IL-8 (D) were measured, with GAPDH as the internal control. Results are based on three independent experiments and shown as mean ± SD. Significant differences are marked with # (pD<D0.001 vs. untreated) and * (pD<D0.001 vs. LPS-only).

### 3.9. Effect of *Rauwolfia serpentina* dry root extracts and reserpine on LPS-induced pro-inflammatory cytokine release in THP 1 cells

LPS stimulation elicited a pronounced inflammatory response in THP-1 monocytes, characterised by robust increases in TNF-α, MCP-1, IL-6, and IL-8 compared with unstimulated controls. Treatment with *Rauwolfia serpentina* root extracts revealed a clear extraction-dependent anti-inflammatory profile, with ethanol-derived fractions consistently outperforming aqueous preparations. TNF-α secretion rose to 54.85 ± 2.19-fold following LPS exposure. Both ethanolic extracts significantly attenuated this response, with greater suppression observed with CEE (27.30 ± 1.35-fold) than with HEE (40.82 ± 1.21-fold). In contrast, HWE and CWE produced minimal reduction. MCP-1 showed the greatest extract sensitivity. While LPS increased MCP-1 to 32.04 ± 1.18-fold, HEE and CEE reduced levels sharply to 6.00 ± 0.55 and 8.31 ± 0.70-fold, respectively. Aqueous extracts failed to confer comparable protection, with MCP-1 remaining elevated. This pattern indicates a strong enrichment of MCP-1-modulating constituents in ethanol-soluble root metabolites. IL-6 followed a similar trend. Ethanolic extracts markedly suppressed IL-6 secretion from the LPS-induced level of 40.78 ± 1.60-fold to 10.66 ± 1.90 (HEE) and 11.40 ± 1.80-fold (CEE), whereas water-based extracts showed little effect. IL-8 displayed comparatively lower responsiveness, with modest reductions observed for ethanolic extracts and negligible changes with aqueous fractions, suggesting the differential sensitivity of this chemokine to crude extract intervention (Figure 7).

**Figure 7.**
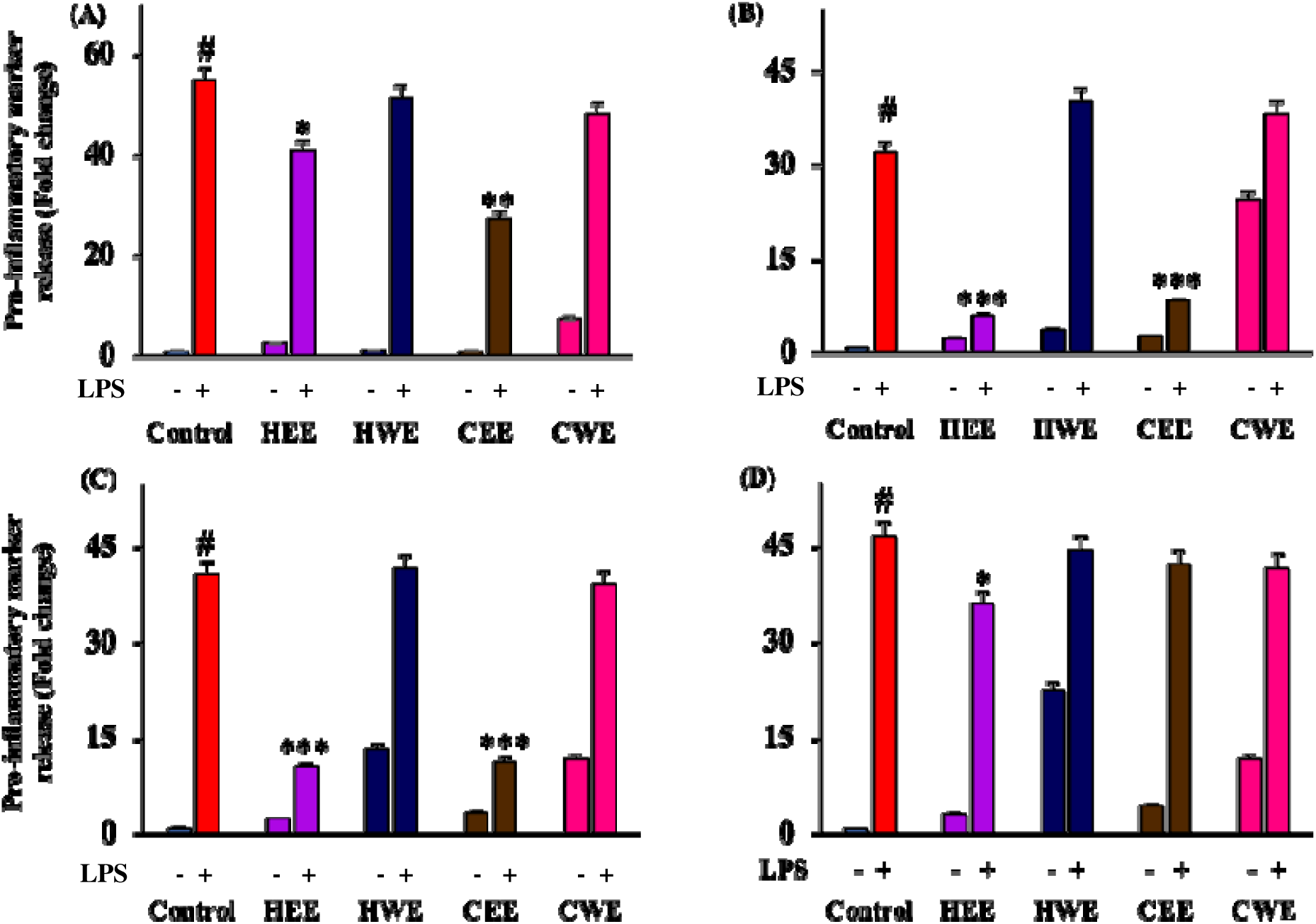
Anti-inflammatory effects of *Rauwolfia serpentina* root extracts in LPS-stimulated THP-1 cells. Cells were pre-treated with *R. serpentina* dry root extracts (120 µg/mL) followed by lipopolysaccharide (LPS) exposure for 3 h. Culture supernatants were collected, and levels of TNF-α (A), MCP-1 (B), IL-6 (C), and IL-8 (D) were quantified by ELISA. Data are expressed as fold change relative to untreated controls and presented as mean ± SD from three independent experiments (n = 3). #p < 0.001 vs. untreated control; *p < 0.001 vs. LPS-treated group.

Reserpine was subsequently evaluated to determine whether this primary alkaloid contributes to the observed anti-inflammatory activity. Reserpine produced a concentration-dependent attenuation of TNF-α and MCP-1, with more substantial effects on IL-6, reducing its expression to 9.75 ± 0.42-fold at 5 µM and 7.39 ± 0.46-fold at 25 µM. IL-8 levels were also reduced, albeit to a lesser extent (Figure 8). Collectively, these findings demonstrate that *Rauwolfia serpentina*, particularly its ethanol-derived fractions, effectively suppresses LPS-induced cytokine release in THP-1 monocytes, with reserpine contributing prominently to the modulation of IL-6 and MCP-1. The data support a biologically meaningful anti-inflammatory action of *R. serpentina,* consistent with its traditional cardiovascular use.

**Figure 8.**
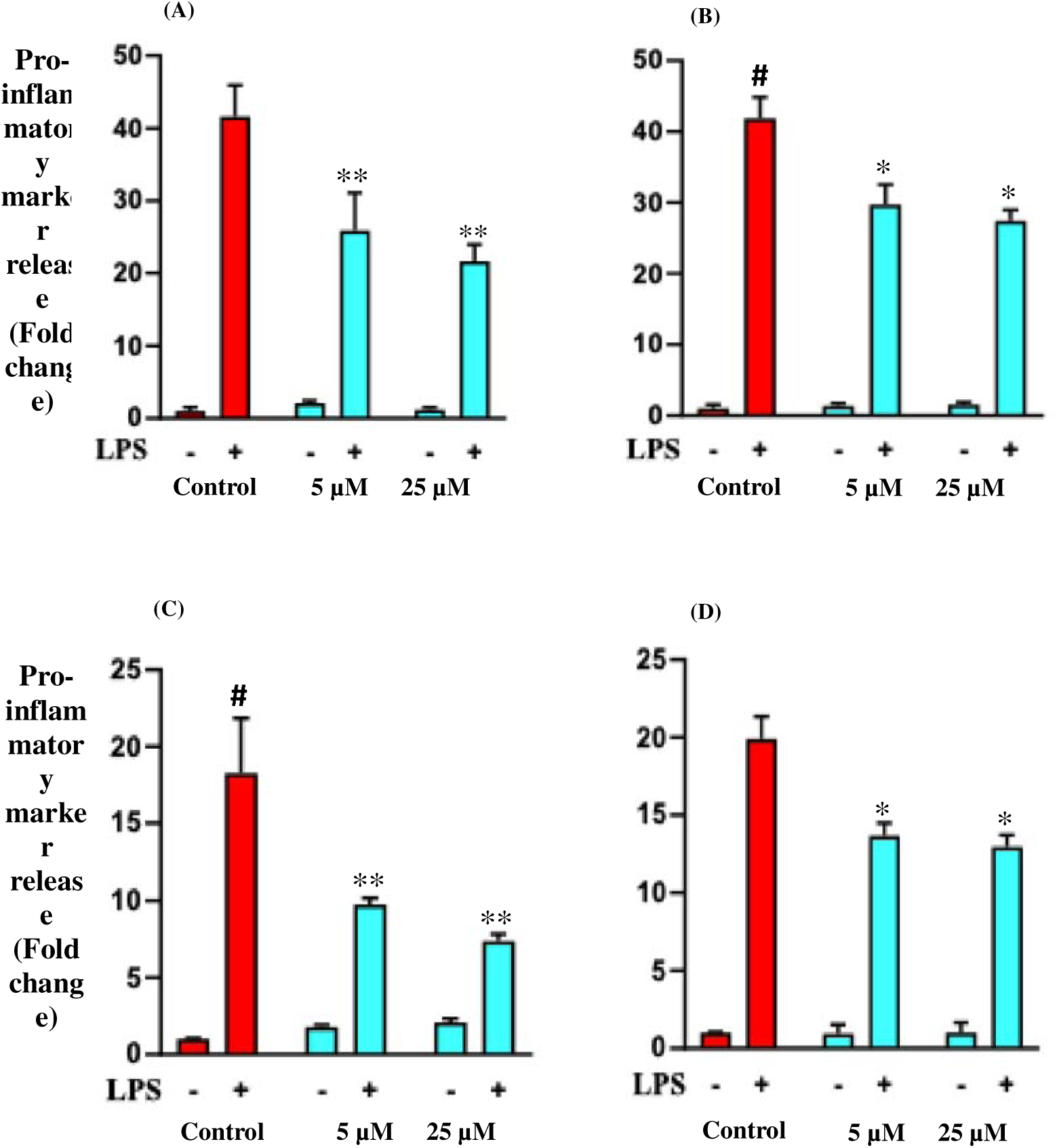
Anti-inflammatory effects of reserpine in LPS-stimulated THP-1 cells. Cells were pre-treated with reserpine (5 and 25 µM) prior to lipopolysaccharide (LPS) stimulation for 3 h. Culture supernatants were collected, and levels of TNF-α (A), MCP-1 (B), IL-6 (C), and IL-8 (D) were quantified by ELISA. Results are expressed as fold change relative to untreated controls and presented as mean ± SD from three independent experiments (n = 3). #p < 0.001 vs. untreated control; *p < 0.001 vs. LPS-treated group.

### 3.10. Molecular docking

In vitro analysis demonstrated that reserpine significantly suppresses the production of inflammatory mediators in THP-1 cells, indicating that modulation of IKKα-dependent NF-κB signalling may underlie the anti-inflammatory activity of *R. serpentina*. To validate this hypothesis, molecular docking analysis was performed using the crystal structure of human IKKα (PDB ID: 5EBZ), a key upstream regulator of NF-κB activation, to assess whether reserpine directly interacts with components of the inflammatory signalling machinery.

Both 5TL and reserpine showed interactions with multiple amino acid residues within the protein kinase domain of IKKα (UniProt ID: O15111; residues 15–302) (Figure 9). Notably, both ligands were also predicted to engage residues within the ATP-binding region (residues 21–29) (Figure 9), suggesting potential interference with kinase catalytic activity.

**Figure 9.**
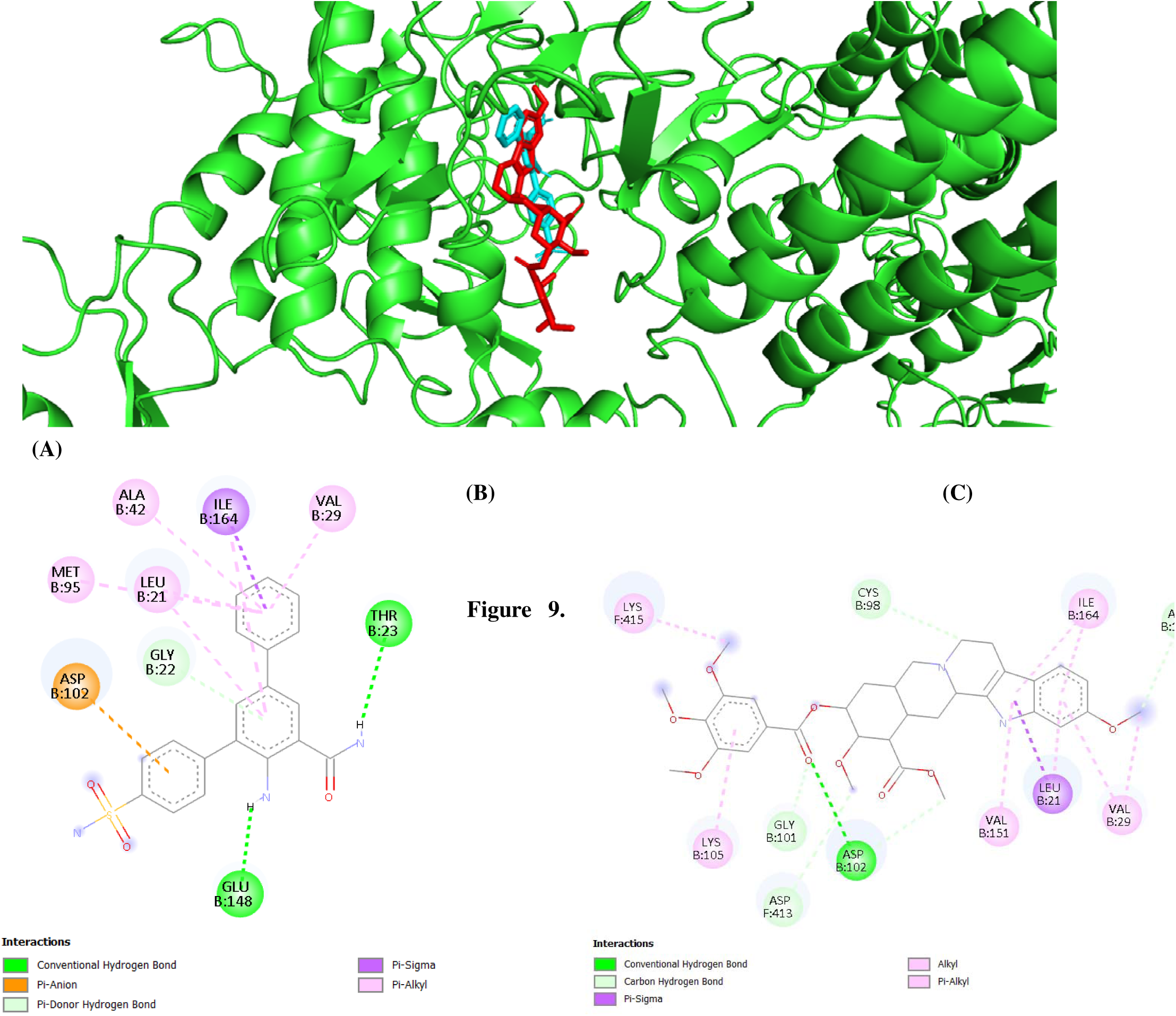
Molecular docking analysis of 5TL and reserpine with human IKKα (PDB ID: 5EBZ). (A) Cartoon representation of the IKKα (green) showing the binding pocket, with docked reserpine (red) and 5TL (cyan). (B) 2D representation of 5TL and IKKα interaction. (C) 2D representation of reserpine and IKKα interaction.

Although the binding affinity of reserpine (–9.4 kcal/mol) was relatively lower than that of 5TL (–10.8 kcal/mol), it still indicates a strong and favourable interaction with IKKα. The localisation of both ligands within the same binding pocket supports the possibility that reserpine may modulate IKKα activity through a mechanism similar to that of 5TL, potentially by competing with ATP binding and thereby inhibiting kinase function.

## 4. Discussion

The present work extends the pharmacological narrative of *Rauwolfia serpentina* beyond its classical antihypertensive identity and situates it within the modern framework of inflammation-driven cardiovascular pathology. Cardiovascular disease is now widely understood as a chronic immunometabolic disorder sustained by oxidative stress and persistent cytokine signalling rather than solely by haemodynamic imbalance (Hotamisligil, 2017; Libby, 2002). Despite this paradigm shift, systematic evaluation of *R. serpentina* in integrated oxidative and inflammatory models has remained limited, with most prior investigations focusing either on isolated alkaloids or on blood pressure outcomes (Lobay, 2015; Shamon and Perez, 2016). The current findings address this gap by demonstrating coordinated antioxidant and anti-inflammatory activity across complementary biochemical, cellular, and computational platforms.

A defining observation is the clear extraction-dependent divergence in biological activity. Ethanolic fractions consistently surpassed aqueous extracts in radical scavenging, inhibition of lipid peroxidation, preservation of catalase activity, and suppression of inflammatory mediators. This pattern is chemically rational. Ethanol preferentially extracts semi-polar indole alkaloids and related secondary metabolites, which constitute the dominant phytochemical class of *R. serpentina* roots (Kumari et al., 2013; Mohammed et al., 2021; Vakil, 1949). Recent phytochemical and metabolomic studies have highlighted the structural diversity of *Rauwolfia* alkaloids and their potential interactions with redox and signalling pathways beyond monoamine depletion (Pathania et al., 2006; Singh et al., 2004). By contrast, aqueous extraction enriches highly polar constituents that may possess lower membrane permeability and reduced interaction with intracellular signalling targets (Dai and Mumper, 2010; Do et al., 2014). The present data therefore reinforce the importance of solvent strategy in botanical standardisation and mechanistic reproducibility.

Among inflammatory mediators, MCP-1 and IL-6 showed the strongest suppression. These cytokines are central regulators of monocyte recruitment, vascular inflammation, and plaque destabilisation (Libby, 2021b; Ridker, 2016). Their selective sensitivity suggests interference with upstream signalling nodes rather than nonspecific antioxidant effects. Although reserpine reproduced a substantial portion of the cytokine modulation, the broader extract activity exceeded that of the isolated compound, supporting phytochemical synergy as recognised in contemporary natural product pharmacology (Williamson, 2001). This distinction is scientifically relevant because most existing Rauwolfia literature reduces biological interpretation to reserpine alone, overlooking potential cooperative interactions among alkaloids and minor metabolites.

The docking analysis further provides mechanistic plausibility. Stable predicted binding of reserpine within the catalytic pocket of IKKα, a kinase central to NF-κB activation (Eberhardt et al., 2021), aligns with the observed transcriptional downregulation of pro-inflammatory genes. While computational, this link bridges phytochemistry with a defined inflammatory signalling axis and complements recent efforts to identify natural inhibitors of IKK-mediated pathways (Valotto Neto et al., 2024; Wadhwa et al., 2022). Such integration of in vitro and in silico evidence remains relatively underexplored for *R. serpentina*, representing a methodological advancement over earlier descriptive studies.

Importantly, this study contributes to an emerging re-evaluation of historically used botanicals within modern immunobiology. By combining metabolomic profiling, oxidative stress models, cytokine analysis, and molecular docking, the work moves beyond symptomatic endpoints and addresses mechanistic dimensions relevant to chronic vascular inflammation. Limitations include reliance on cell-based and *ex vivo* systems, absence of targeted alkaloid quantification, and lack of *in vivo* validation. Nevertheless, the convergence of redox modulation, cytokine suppression, and predicted kinase engagement provides a coherent mechanistic framework that advances current understanding and positions *R. serpentina* as a multi-target botanical candidate for inflammation-driven cardiovascular complications.

## 5. Conclusions

This study demonstrates that *R. serpentina* root extracts, particularly ethanol-derived extracts, exert coordinated antioxidant and anti-inflammatory effects by modulating oxidative stress, inflammatory mediators, and key signalling pathways. The findings highlight extraction chemistry as a critical determinant of bioactivity and support a synergistic contribution of indole alkaloids rather than reliance on reserpine alone. Collectively, the results strengthen the scientific basis for *R. serpentina* as a complementary botanical candidate for managing inflammation-associated cardiovascular complications and provide a focused platform for future translational and in vivo investigations.

## 6. Acknowledgement

The author sincerely acknowledges the late Prof. Sarada D. Tetali for her guidance and support throughout the research period. Financial support from the University of Hyderabad - Institution of Eminence (UoH-IoE) (Grant No. UoH-IoE-RC3-21-027).

