## Supplementary Figures for "Multimodal antioxidant and anti-inflammatory activity of *Rauwolfia serpentina* root extracts in experimental models"

% DPPH Scavenging Activity

| Standard compound (µg/mL) | abs at 517nm |
| --- | --- |
| 0 | 0 |
| 1 | 15.0342 |
| 2 | 29.4421 |
| 3 | 35.8969 |
| 4 | 47.637 |
| 5 | 57.8542 |
| 6 | 65.2142 |

| Standard compound (µg/mL) | GA conc in µg/ml |
| --- | --- |
| 0 | 0 |
| 0.5 | 7.52161 |
| 1 | 22.2044 |
| 1.5 | 31.9868 |
| 2 | 44.4621 |
| 2.5 | 57.8027 |
| 3 | 68.0925 |
| 3.5 | 75.0472 |

Ascorbic acid

Gallic acid

**Supplementary Figure 1.** Free radical inhibition measured via DPPH was carried out for standard antioxidant compounds, gallic acid and ascorbic acid. The results are expressed as mean ± standard deviation, based on triplicate measurements (n = 3). All data points showed statistically significant differences, with p < 0.001.

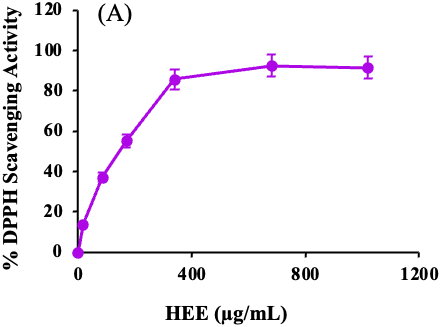

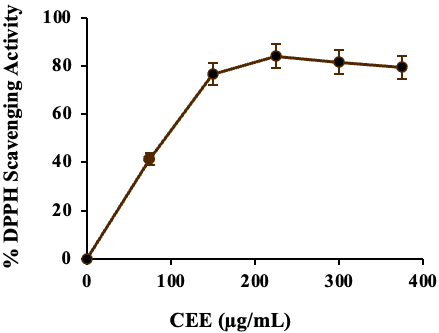

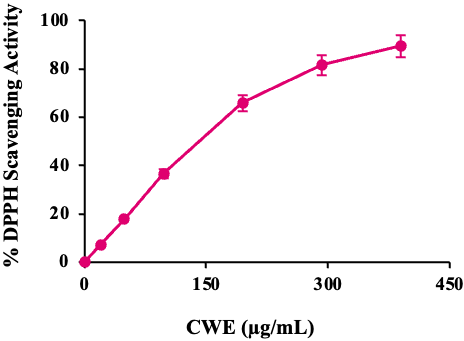

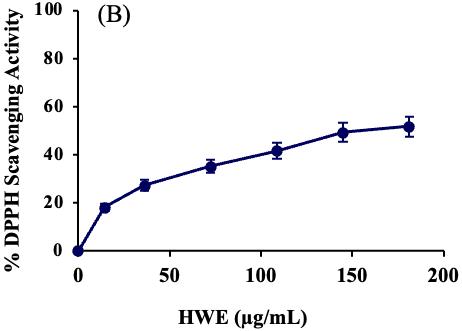

(C)

(D)

% DPPH Scavenging Activity

| HEE (µg/mL) | % of inhbition |
| --- | --- |
| 0 | 0 |
| 17 | 13.8495 |
| 85 | 37.1283 |
| 170 | 55.1813 |
| 340 | 85.6705 |
| 680 | 92.4602 |
| 1020 | 91.3825 |

% DPPH Scavenging Activity

| CEE (µg/mL) | IC50 |
| --- | --- |
| 0 | 0 |
| 75 | 41.2552 |
| 150 | 76.5798 |
| 225 | 83.9501 |
| 300 | 81.5204 |
| 375 | 79.4682 |

% DPPH Scavenging Activity

| CWE (µg/mL) | % of inhbition |
| --- | --- |
| 0 | 0 |
| 19.5 | 7.2013 |
| 48.75 | 17.8053 |
| 97.5 | 36.6249 |
| 195 | 65.902 |
| 292.5 | 81.6344 |
| 390 | 89.5033 |

% DPPH Scavenging Activity

| HWE (µg/mL) | % of inhbition |
| --- | --- |
| 0 | 0 |
| 14.5 | 18.0091 |
| 36.25 | 27.2353 |
| 72.5 | 35.1207 |
| 108.75 | 41.6427 |
| 145 | 49.2021 |
| 181.25 | 51.7282 |

**Supplementary Figure 2.** Free radical inhibition measured via DPPH was carried out for standard antioxidant compounds (A), along with the cold ethanolic extract (CEE) (B), hot water extract (HWE) (C), and cold water extract (CWE) (D) of *Piper nigrum* dried fruits. The results are expressed as mean ± standard deviation, based on triplicate measurements (n = 3). All data points showed statistically significant differences, with p < 0.001.

% DPPH Scavenging Activity

| Reserpine (µM) | % of inhbition |
| --- | --- |
| 0 | 0 |
| 2.5 | 17.2326 |
| 5 | 21.9207 |
| 10 | 27 |
| 20 | 35.9694 |
| 30 | 41.9323 |
| 40 | 47.3053 |
| 50 | 50.5752 |
| 75 | 50.4942 |

**Supplementary Figure 3.** DPPH radical scavenging studies: piperine. The results are expressed as mean ± standard deviation, based on triplicate measurements (n = 3). All data points showed statistically significant differences, with p < 0.001.

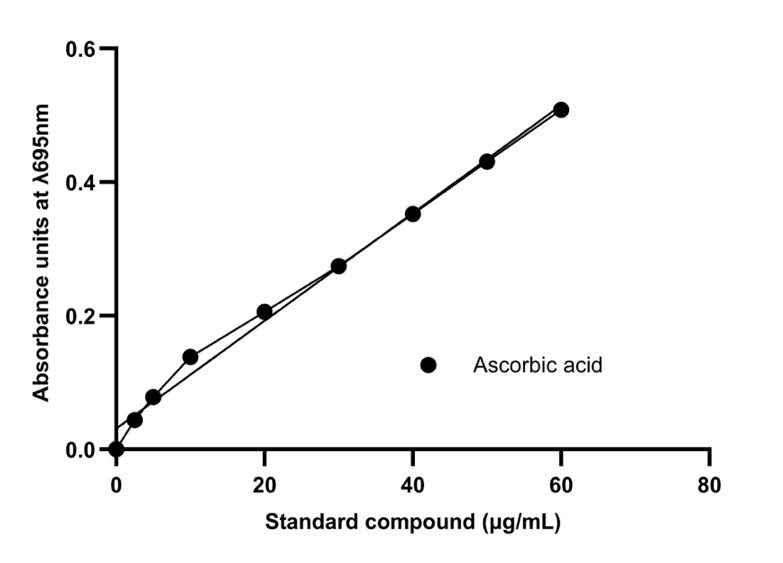

**(A)**

**(B)**

|  | Total antioxidant capacity (mg GAE/g dw) |
| --- | --- |
| HEE | 10.81 |
| HWE | 16.86 |
| CEE | 10.36 |
| CWE | 10.46 |

**Supplementary Figure 4.** (A) Standard curve for total antioxidant capacity, (B) Total antioxidant capacity of different extracts of *Rauwolfia serpentina*

(C)

(B)

(A)

Total terpenoid content (mg LE/g dw)

|  | Total terpenoid content (mg LE/g) |
| --- | --- |
| CEE | 2.19 |
| HEE | 2.78 |
| CWE | 0.32 |
| HWE | 0.56 |

Total flavonoid content (mg QE/g dw)

|  | Total flavonoid content (mg QE/g) |
| --- | --- |
| CEE | 89.93 |
| HEE | 59.67 |
| CWE | 14.15 |
| HWE | 15.57 |

Total phenol content (mg GAE/g dw)

|  | Total phenol content (mg GAE/g) |
| --- | --- |
| CEE | 67.66 |
| HEE | 24.07 |
| CWE | 25.57 |
| HWE | 28.92 |

**Supplementary Figure 5.** Total Phenol (A), Flavonoid (B), Terpenoid(C) content of *R. serpentina* dry root extracts. The data, based on a sample size of three, include standard deviation values, and all measurements were found to be statistically significant at a confidence level of p < 0.001.

| R. serpentina dry root extracts (160 dwt µg/mL) | % cell viability |
| --- | --- |
| Control | 100 |
| HEE | 99 |
| HWE | 97 |
| CEE | 97 |
| CWE | 96 |

| Reserpine (µM) | % cell viability |
| --- | --- |
| Control | 100 |
| 5 | 98 |
| 25 | 95 |
| 50 | 97 |
| 100 | 88 |
| 150 | 83 |
| 200 | 77 |

**Supplementary Figure 6.** To evaluate cytotoxicity, the viability of THP-1 cells was examined using the MTT assay following treatment with both aqueous and alcoholic extracts of *Rauwolfia serpentina* and Reserpine. Cells were incubated overnight at 37°C with 5% CO₂ in the presence of 160 ± 5 µg/mL of *R. serpentina* extract and reserpine in the range of 5 to 250 µM. Control wells contained only media combined with the respective extract concentrations and metabolite, but without any cells. All values reflect the average ± standard deviation from three separate experimental replicates.

(A)

(B)

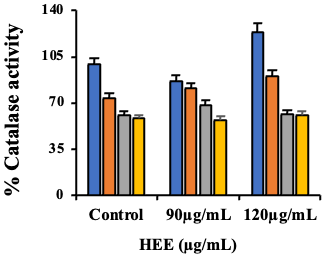

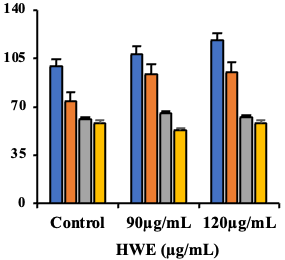

(C)

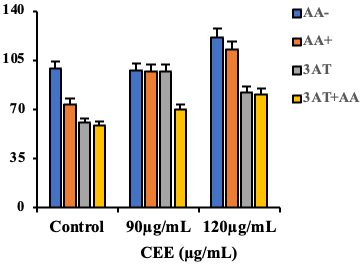

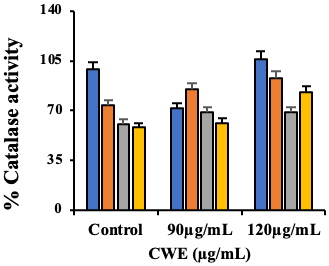

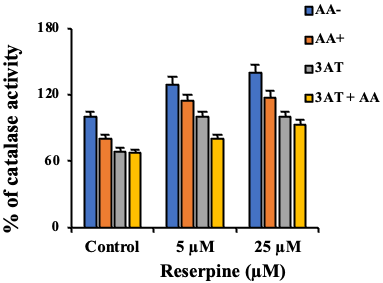

(D)

(E)

**Supplementary Figure 7.** Effect of R. serpenitna extracts on catalase activity (A) piperine (B) reserpine (C) ellagic acid on Catalase activity. The results are expressed as mean ± standard deviation, based on triplicate measurements (n = 3). All data points showed statistically significant differences, with p < 0.001.

% Catalase activity

| HEE (µg/mL) | AA- | AA+ | 3AT | 3AT+AA |
| --- | --- | --- | --- | --- |
| Control | 99.32 | 73.78 | 60.74 | 58.26 |
| 90µg/mL | 86.93 | 81.07 | 68.57 | 57.35 |
| 120µg/mL | 123.99 | 90.53 | 61.41 | 60.71 |

| HWE (µg/mL) | AA- | AA+ | 3AT | 3AT+AA |
| --- | --- | --- | --- | --- |
| Control | 99.32 | 73.78 | 60.74 | 58.26 |
| 90µg/mL | 108.36 | 93.65 | 65.51 | 52.81 |
| 120µg/mL | 118.18 | 95.4 | 62.5 | 58.41 |

| CEE (µg/mL) | AA- | AA+ | 3AT | 3AT+AA |
| --- | --- | --- | --- | --- |
| Control | 99.32 | 73.78 | 60.74 | 58.26 |
| 90µg/mL | 98.06 | 96.98 | 97.09 | 69.84 |
| 120µg/mL | 121.37 | 112.89 | 82.3 | 80.93 |

% Catalase activity

| CWE (µg/mL) | AA- | AA+ | 3AT | 3AT+AA |
| --- | --- | --- | --- | --- |
| Control | 99.32 | 73.78 | 60.74 | 58.26 |
| 90µg/mL | 71.78 | 85.27 | 68.87 | 61.33 |
| 120µg/mL | 106.2 | 92.73 | 68.87 | 82.99 |

% of catalase activity

| Reserpine (µM) | AA- | AA+ | 3AT | 3AT + AA |
| --- | --- | --- | --- | --- |
| Control | 100 | 80.19 | 68.68 | 67.47 |
| 5 µM | 129.76 | 114.8 | 100.31 | 80.2 |
| 25 µM | 140.58 | 117.99 | 100.01 | 93.19 |
