## Supplementary Table 1 for "Multimodal antioxidant and anti-inflammatory activity of *Rauwolfia serpentina* root extracts in experimental models"

**Supplementary Table 1*. Rauwolfia serpentina* and other *Rauwolfia* species-specific secondary metabolites from the current LC-MS investigation of *R. serpentina* dried root extracts.**

| **Si No** | **Molecular formula** | **RT** | **Class of compounds** | **Molecular formula** | **Abundance** | **Plant name** | **Plant part** |
| --- | --- | --- | --- | --- | --- | --- | --- |
| 1 | (16S)-Sarpagan-17-ol | 17.09 | Alkaloid | C_19_H_22_N_2_O | 53650.44 | *Rauwolfia densiflora* | Leaves |
| 2 | 17-O-Acetylajmaline | 17.68 | Alkaloid | C₂₂H₂₈N₂O₃ | 16100000.0 | *Rauwolfia serpentina* | Root bark |
| 3 | 3-alpha(S)-Strictosidine | 15.95 | Alkaloid | C₂₇H₃₄N₂O₉ | 127726.6 | *Rauwolfia serpentina* | Roots |
| 4 | 3-Oxorhazinilam | 1.21 | Alkaloid | C₁₉H₂₀N₂O₂ | 219289.6 | *Rauwolfia serpentina* | Cell Culture |
| 5 | 3,4,5-Trimethoxybenzoic acid | 6.41 | Phenol | C₁₀H₁₂O₅ | 141357.6 | *Rauwolfia serpentina* | Roots |
| 6 | 7-Dehydrositosterol | 26.02 | Phytosterol | C₂₉H₄₈O | 63650.1 | *Rauwolfia serpentina* | Roots |
| 7 | 12-Hydroxyajmaline | 13.58 | Alkaloid | C_20_H_26_N_2_O_3_ | 7008828.8 | *Rauwolfia serpentina* | Cell Culture |
| 8 | Ajmalicine | 17.30 | Alkaloid | C₂₁H₂₄N₂O₃ | 15900000.0 | *Rauwolfia serpentina* | Roots |
| 9 | Ajmalidine | 27.38 | Alkaloid | C₂₀H₂₄N₂O₂ | 184901.4 | *Rauwolfia serpentina* | Root bark |
| 10 | Ajmaline | 7.63 | Alkaloid | C₂₀H₂₆N₂O₂ | 2771447.0 | *Rauwolfia serpentina* | Roots |
| 11 | Ajmalinimine | 21.20 | Alkaloid | C₂₄H₃₀N₂O₄ | 723712.4 | *Rauwolfia serpentina* | Roots |
| 12 | Alstonine | 15.42 | Alkaloid | C₂₁H₂₀N₂O₃ | 98147.6 | *Rauwolfia serpentina* | Roots |
| 13 | Arachidic acid | 27.42 | Fatty acid | C_20_H_40_O_2_ | 220560.4 | *Rauwolfia serpentina* | seed oils |
| 14 | Arbutin | 4.76 | Phenolic glycoside | C₁₂H₁₆O₇ | 137752.4 | *Rauwolfia serpentina* | Suspension cell culture |
| 15 | Aricine | 2.61 | Alkaloid | C₂₂H₂₆N₂O₄ | 200779.6 | *Rauwolfia serpentina* | Roots |
| 16 | Aricine | 2.61 | Alkaloid | C₂₂H₂₆N₂O₄ | 200779.6 | *Rauwolfia volkensii* | Roots |
| 17 | beta-Sitosterol | 27.41 | Phytosterol | C₂₉H₅₀O | 42051.4 | *Rauwolfia serpentina* | Roots |
| 18 | Deserpidine | 23.98 | Alkaloid | C₃₂H₃₈N₂O₈ | 56787.3 | *Rauwolfia canescens* | Leaves |
| 19 | Deserpidine | 23.98 | Alkaloid | C₃₂H₃₈N₂O₈ | 56787.3 | *Rauwolfia serpentina* | Roots |
| 20 | Docosanoic acid | 12.13 | Fatty acid | C₂₂H₄₄O₂ | 210572.5 | *Rauwolfia serpentina* | Leaves |
| 21 | gamma-Sitosterol | 33.07 | Phytosterol | C₂₉H₅₀O | 15923.0 | *Rauwolfia serpentina* | Seeds |
| 22 | Geissoschizol | 23.87 | Alkaloid | C₁₉H₂₄N₂O | 126135.8 | *Rauwolfia serpentina* | Roots |
| 23 | Glomeratose A | 29.06 | Phenylpropanoid | C₂₄H₃₄O₁₅ | 74478.7 | *Rauwolfia serpentina* | Root bark |
| 24 | Isoajmaline | 8.71 | Alkaloid | C₂₀H₂₆N₂O₂ | 2513479.3 | *Rauwolfia serpentina* | Roots |
| 25 | Isoreserpiline | 6.24 | Alkaloid | C₂₃H₂₈N₂O₅ | 37727.9 | *Rauwolfia cambodiania* | Roots |
| 26 | Isosandwicine | 9.06 | Alkaloid | C₂₀H₂₆N₂O₂ | 466669.3 | *Rauwolfia serpentina* | Cell culture |
| 27 | Lauric acid | 7.452 | Fatty acid | C_12_H_24_O_2_ | 408923.406 | *Rauwolfia serpentina* | Seed oil |
| 28 | Loganic acid | 33.43 | Terpenoid | C₁₆H₂₄O₁₀ | 25378.4 | *Rauwolfia serpentina* | Roots |
| 29 | Loganin | 19.14 | Terpenoid | C₁₇H₂₆O₁₀ | 400709.3 | *Rauwolfia grandiflora* | Bark |
| 30 | Macrophylline | 27.51 | Alkaloid | C₁₃H₂₁NO₃ | 1207181.8 | *Rauwolfia caffra* | Stems |
| 31 | Myristic acid | 34.58 | Fatty acid | C_14_H_28_O_2_ | 108479.674 | *Rauwolfia serpentina* | Seed oil |
| 32 | Norajmaline | 10.42 | Alkaloid | C₁₉H₂₄N₂O₂ | 1850671.5 | *Rauwolfia serpentina* | Roots |
| 33 | Normacusine B | 4.69 | Alkaloid | C₁₉H₂₂N₂O | 45736.2 | *Rauwolfia serpentina* | Roots |
| 34 | Oleic acid | 27.49 | Fatty acid | C₁₈H₃₄O₂ | 103884.2 | *Rauwolfia serpentina* | Roots |
| 35 | Papaverine | 14.90 | Alkaloid | C₂₀H₂₁NO₄ | 4044601.1 | *Rauwolfia serpentina* | Roots |
| 36 | Palmitic acid | 1.944 | Fatty acid | C_16_H_32_O_2_ | 183603.2 | *Rauwolfia serpentina* | Seed oil |
| 37 | Raucaffricine | 26.95 | Alkaloid | C₂₂H₂₈N₂O₃ | 199749.3 | *Rauwolfia serpentina* | Roots |
| 38 | Raucaffrinoline | 17.72 | Alkaloid | C₂₇H₃₄N₂O₉ | 3136715.5 | *Rauwolfia serpentina* | Root culture |
| 39 | Raumacline | 18.88 | Alkaloid | C₁₉H₂₀N₂O₂ | 305713.7 | *Rauwolfia serpentina* | Root culture |
| 40 | Raunescine | 32.42 | Alkaloid | C₁₀H₁₂O₅ | 77102.1 | *Rauwolfia serpentina* | Roots |
| 41 | Raupine (Sarpagine) | 8.91 | Alkaloid | C₂₉H₄₈O | 130739.4 | *Rauwolfia serpentina* | Roots |
| 42 | rauwolscine | 29.96 | Alkaloid | C₂₅H₃₄O₁₄ | 35232.4 | *Rauwolfia serpentina* | Roots |
| 43 | Rescinnamidine | 9.18 | Alkaloid | C₂₁H₂₄N₂O₃ | 1588942.8 | *Rauwolfia serpentina* | Roots |
| 44 | Rescinnamine | 18.35 | Alkaloid | C_33_H_40_N_2_O_9_ | 90402.9 | *Rauwolfia serpentina* | Roots |
| 45 | Reserpiline | 34.80 | Alkaloid | C₂₄H₃₀N₂O₄ | 9135.9 | *Rauwolfia serpentina* | Stems |
| 46 | Reserpine N-Oxide | 43.02 | Alkaloid | C₂₁H₂₀N₂O₃ | 86319.3 | *Rauwolfia serpentina* | Roots |
| 47 | Rhazinilam | 3.69 | Alkaloid | C₁₉H₂₂N₂O | 83895.6 | *Rauwolfia serpentina* | Hybrid cell culture |
| 48 | Sarpagine | 6.86 | Alkaloid | C₁₂H₁₆O₇ | 176274.8 | *Rauwolfia serpentina* | Roots |
| 49 | Secologanin | 32.76 | Terpenoid | C₂₂H₂₆N₂O₄ | 88622.4 | *Rauwolfia serpentina* | Suspension culture |
| 50 | Serpentine | 7.09 | Alkaloid | C₂₂H₂₆N₂O₄ | 187312.2 | *Rauwolfia serpentina* | Roots |
| 51 | Serpentinine | 18.18 | Alkaloid | C₂₉H₅₀O | 685.4 | *Rauwolfia serpentina* | Roots |
| 52 | Stearic acid | 4.30 | Fatty acid | C_18_H_36_O_2_ | 137728.2 | *Rauwolfia serpentina* | Seeds |
| 53 | Stemmadenine | 14.24 | Alkaloid | C_21_H_26_N_2_O_3_ | 9647018.3 | *Rauvolfia serpentina × Rhazya*  *stricta* | Cell culture |
| 54 | Stemmadenine | 14.24 | Alkaloid | C₃₂H₃₈N₂O₈ | 9647018.2 | *Rauwolfia serpentina x Rhazya stricta* | Leaves |
| 55 | Stigmasterol | 14.24 | Phytosterol | C_29_H_48_O | 140740.7 | *Rauwolfia serpentina* | Roots |
| 56 | Swertiaside | 33.05 | Terpenoid | C_23_H₂_8_O₁_2_ | 67480.6 | *Rauwolfia serpentina* | Roots |
| 57 | Tetrahydroalstonine | 16.88 | Alkaloid | C_21_H_24_N_2_O_3_ | 2281636.0 | *Rauwolfia serpentina* | Roots |
| 58 | Tetraphyllicine | 25.78 | Alkaloid | C₂₀H₂₆N₂O₂ | 170990.1 | *Rauwolfia serpentina* | Roots |
| 59 | Tetraphylline | 36.09 | Alkaloid | C_22_H_26_N_2_O_4_ | 6871.4 | *Rauwolfia serpentina* | Root bark |
| 60 | Thebaine | 10.60 | Alkaloid | C_19_H_21_NO_3_ | 9500729.6 | *Rauwolfia serpentina* | Roots |
| 61 | Tryptamine | 0.81 | Alkaloid | C_10_H_12_N_2_ | 2039033.2 | *Rauwolfia serpentina* | Cell culture |
| 62 | Tubotaiwine | 5.56 | Alkaloid | C_20_H_24_N_2_O_2_ | 235189.4 | *Rauvolfia serpentina × Rhazya stricta* | Cell culture |
| 63 | Vellosimine | 1.25 | Alkaloid | C_19_H_20_N_2_O | 928880.5 | *Rauwolfia serpentina* | Cell culture |
| 64 | Vinburnine | 1.17 | Alkaloid | C₁₉H₂_2_N₂O | 970905.3 | *Rauwolfia serpentina* | Cell culture |
| 65 | Vinorine | 5.60 | Alkaloid | C_21_H_22_N_2_O_2_ | 133848.7 | *Rauwolfia serpentina* | Roots |
| 66 | Vomilenine | 16.83 | Alkaloid | C_21_H_22_N_2_O_3_ | 12700000.0 | *Rauwolfia serpentina* | Roots |
| 67 | Yohimbic Acid | 37.13 | Alkaloid | C_20_H_24_N_2_O_3_ | 22662.6 | *Rauwolfia serpentina* | Roots |
| 68 | Yohimbine | 13.53 | Alkaloid | C_21_H_26_N_2_O_3_ | 17700000.0 | *Rauwolfia serpentina* | Roots |
